# Mice remember experiences via conspecific-context: models of social episodic-like memory

**DOI:** 10.1101/2023.08.04.552022

**Authors:** T. W. Ross, S. L. Poulter, C. Lever, A. Easton

## Abstract

The ability to remember unique past events (episodic memory) may be an evolutionarily conserved function, with accumulating evidence of episodic-(like) memory processing in rodents. In humans, it likely contributes to successful complex social networking. Rodents, arguably the most used laboratory models, are also rather social animals. However, many behavioural paradigms are devoid of sociality, and commonly-used social spontaneous recognition tasks (SRTs) are open to non-episodic strategies based upon familiarity. We address this gap by developing new SRT variants. Here, in object-in-context SRTs, we asked if context could be specified by the presence/absence of either a conspecific (experiment 1) or an additional local object (experiment 2). We show that mice readily used the conspecific as contextual information to distinguish unique episodes in memory. In contrast, no coherent behavioural response emerged when an additional object was used as a potential context specifier. Further, in a new social conspecific-in-context SRT (experiment 3) where environment-based change was the context specifier, mice preferably explored a more recently-seen familiar conspecific associated with contextual mismatch, over a less recently-seen familiar conspecific presented in the same context. The results argue that, in incidental SRT conditions, mice readily incorporate conspecific cue information into episodic-like memory. Thus, the tasks offer different ways to assess and further understand the mechanisms at work in social episodic-like memory processing.

---

Many animals are innately social species and live in groups^1, 2^. The demand (upon individuals) of maintaining complex social dynamics within group living, is thought to have contributed to evolutionary shaping of the brain^1, 2^. Recognition memory is a necessary cognitive capacity to enable successful complex social living and networking^3–6^. It can be modelled as a dual process where familiarity (knowing) is distinct from recollection (remembering)^7, 8^. You may recognise that a conspecific is familiar, but you may not remember any experiences of how you may know them. This remembering, a core feature of episodic memory, one’s memory for unique past events^9^, allows for the basis of more complex sociality^3, 4^. For example, being vigilant of a once trustworthy conspecific that you deem is no longer trustworthy, because you remember the occasion that they stole your family’s share of food (see^3, 10^).

When considering the evolutionary trajectory of episodic memory, some argue that in its essence it is a human specific ability^4, 11^. Alternatively, some argue that a form of episodic memory exists in many species, evidenced behaviourally^12–14^ and by evolutionarily conserved neural mechanisms^15, 16^. Hence, a more nuanced approach seeks to understand what elements of episodic memory are shared between species^17^. In this way, episodic-like memory has been behaviourally characterised as memory for a simultaneous integration of content (what) in its specific spatial arrangement (where) and temporal context (when)^12^. However, ‘when’ is not the only way to specify episodes in memory. Animals may struggle to remember episodes via an absolute moment in time and may instead rely upon ‘how long ago’^18, 19^ (recency-based memory - susceptible to familiarity processing^14, 20, 21^). Thus, integrated what-where stimuli can also be remembered via contextual specifiers, including the physical environment, acting as an ‘occasion setter’^14^. This is a more holistic interpretation which includes (but is not limited to) ‘when’ being used as the episode specifier. In this work, we further explore temporal cues and the role of contextual specifiers beyond the physical environment.

Rodents have been seen to display episodic-like memory in spontaneous object recognition (SOR) paradigms using both temporal context and other contextual markers to specify and remember episodes^24–26^. Interestingly, however, where recency-based ‘when’ and context-based recognition strategies are available in the same tasks^27^, it seems to be context-based strategies that more commonly shape behaviour overall^28–30^. This raises the question of what kinds of information can readily be used as contextual specifiers, enough to motivationally drive behavioural output during retrieval (over recency-based strategies or randomness), especially in such ‘spontaneous’ tasks where there are minimal explicit external reinforcers being used by experimenters^31^.

In nearly all context-SOR rodent studies, changes in context are operationalised as discrete manipulations of the global environment, typically involving changes in visuo-tactile and or geometric cue information (of walls, floors, and the extra-maze)^24, 28, 30, 32^. It is well-established that these kinds of changes can evoke profound changes in ensembles of hippocampal neurons (see^33, 34^), and such ensemble coding changes are thought to contribute to contextual episodic-like memory processing^16, 35–38^. Yet, even in an experimental setting rodents can naturally form complex social networks^39^, can learn and retrieve hierarchal social status information^40^ and display pro-social behaviour dependent on nurture factors^41^. Thus, such work suggests that rodents may flexibly incorporate social information into episodic-like memory (c.f.^42^).

Here, we use two new variants of the object-in-context SOR paradigm. In the first object-in-context SOR experiment (Experiment 1), we asked if ‘context’ could be specified via the presence/absence of a freely roaming conspecific (**Fig. 1**). In a second object-in-context SOR experiment (Experiment 2), we asked if ‘context’ could be specified via the presence/absence of an additional static local object. In the first experiment, we show that mice readily use conspecific presence and absence as contextual information to separate and distinguish particular events, and this episodic-like strategy was over an object recency-based strategy that was also possible in the SOR. In the second object-in-context SOR experiment, the presence and absence of an additional local object (kept the same throughout the testing session) did not elicit a coherent recognition strategy.

**Figure 1.**
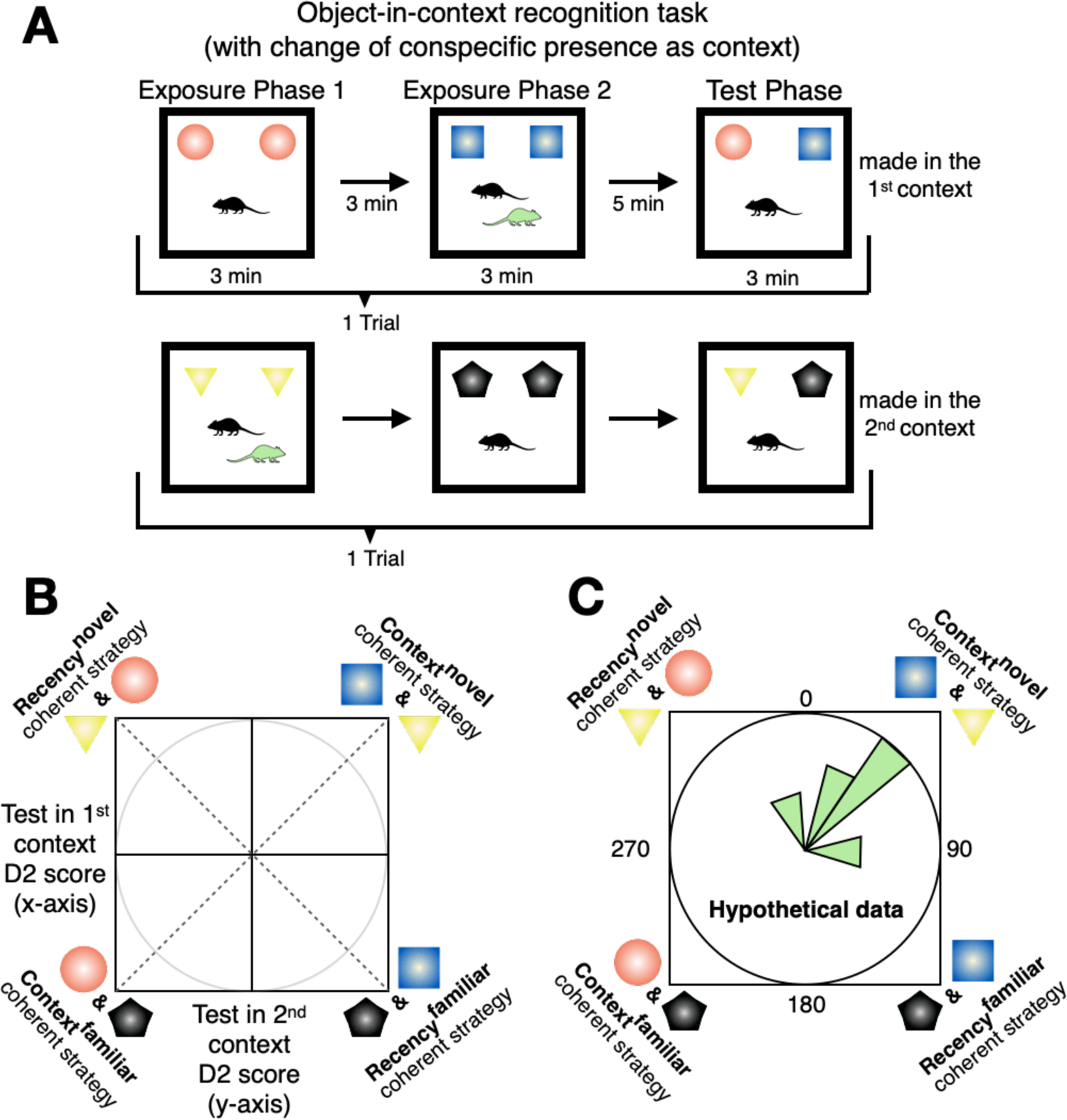
Schematics of the object-in-context recognition task with a conspecific partner as context example. (**A**) In the object-in-context task mice are presented with 2 exposure phases, both containing the same objects. ‘Context’ was specified as presence/absence of a freely roaming conspecific (experiment 1; green mouse in A; same-sex cage and litter mates). The test phase contained a copy of each object experienced from the exposure phases and was only made when the experimental subject was alone (black mouse in A; see main text for reasoning). The test phase could be made in the 1^st^ context (upper), or it could be made in the 2^nd^ context (lower). Thus, two exposure phases and a test phase constituted a single trial (4 trials in each experimental session per animal). (**B**) The D2 ratio scores from test in the 1^st^ context trials can be plotted against test in the 2^nd^ context trial D2 scores and expressed as circular data (via an arctangent function) to test for potentially coherent behavioural strategies across the two types of trials (see^27^ and methods). Context^novel^ and recency^novel^ denote exploration based on object novelty preference, whereas context^familiar^ and recency^familiar^ denote familiarity-based exploratory preference. (**C**) Depicts hypothetical circular data plotted in an angular histogram. In this hypothetical example, the circular mean is ∼45°, suggesting common context^novel^ strategy. One can conduct inferential circular statistics asking whether the data is uniformly distributed around the circle or not. In this hypothetical example, if the data is not uniformly distributed around the circle, and thus significantly clustered around the mean of ∼45°, this is indicative of a coherent context^novel^ strategy. Such circular analyses are contingent upon evidence of behavioural recognition preference differing to chance level performance and can enhance explanatory power of the spontaneous recognition task data in terms of strategy.

We also developed a new conspecific-in-context SR task (Experiment 3, **Fig. 2**), based broadly on the model of the standard object-in-context SR task, but employing conspecifics instead of objects. Thus: a) as per the usual convention, context was specified via environment-based change of the floor and wall visuo-tactile cues; b) conspecifics were kept in stable locations (like objects) within wire cups. Here, just as with the standard object-in-context SR task, we asked if mice could detect, and thus preferentially explore, a novel conspecific-in-context configuration (mismatch) over a previously presented conspecific-in-context configuration. To directly pit contextual mismatch against recency-based exploration, we introduced a third conspecific in the second exposure phase, so that in the test phase the conspecific who was not part of the contextual mismatch was seen longer ago (**Fig 2**). In this way, two novelty-oriented discriminatory strategies were available: 1) explore the conspecific more in the novel conspecific-in-context configuration (context mismatch strategy); 2) explore the conspecific more who was seen longer ago (recency strategy). As we shall see, the results favoured a conspecific-in-context episodic-like memory account.

**Figure 2.**
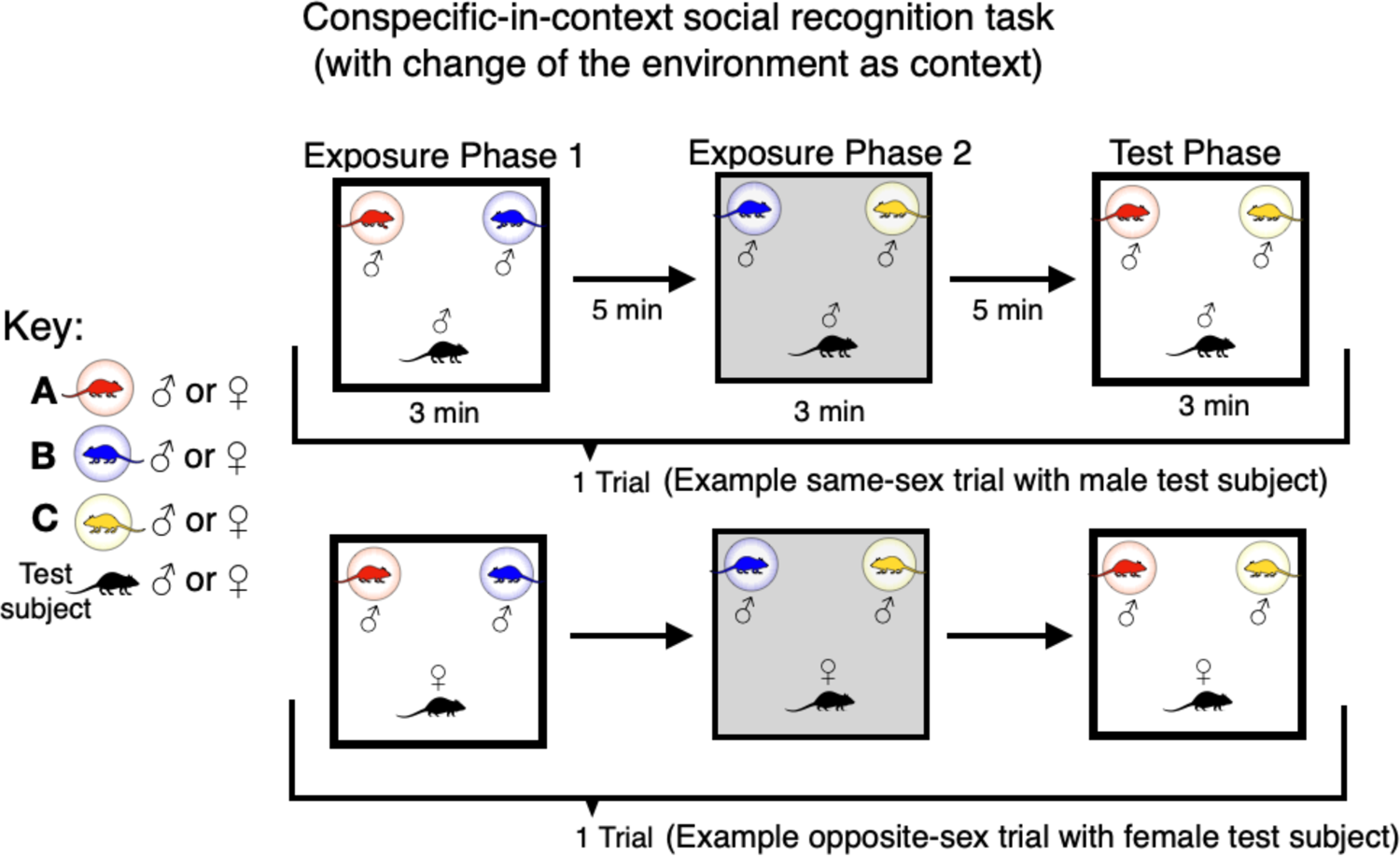
Schematics of the social conspecific-in-context recognition task variant with environment-based change as the context specifier. The conspecific-in-context task was constituted by 3 phases forming a single trial (2 exposure phases, left and middle, and a test phase, right). Here, context was specified via change of the physical environment. In the test phase, both conspecifics should be familiar (based on experience from the exposure phases), but one was seen less recently presented in the same context and place (A; red) and the other was more recently seen, yet now presented in a contextual mismatch (C; yellow). Conspecifics were same-sex cage and littermates in two separate sessions of a single trial per animal (an example male same-sex trial is shown; upper). However, in a final session we also tested opposite-sex littermates (an example opposite-sex trial with a female test subject is shown; lower).

## Results

### Experiment 1: conspecific presence/absence is sufficient to act as a contextual specifier for mice to remember episodes

In this object-in-context task SOR variant (**Fig. 1A**), context was specified by the presence and absence of a freely roaming conspecific partner for experiment 1. This partner was a same-sex littermate and cagemate of the subject. We tested 10 subjects. Subject-partner dyads were kept the same throughout the testing session (a session consisted of 4 trials) and so was the experimental environment. We have recently shown that in object-in-context SOR tasks, animals can either use a recency-based strategy (ignorant of contextual information) or a context-dependent strategy, where the novelty stems from the contextual mismatch at test^27^. The test phase can be situated in the 1^st^ context or in the 2^nd^ context. We ran tests in both contexts for each mouse, and it was imperative to analyse both types of trials separately to assess overall coherent recognition behavioural strategy^27, 43^ (**Fig. 1**).

It was important to impose a restriction upon the test phase; namely, that the experimental subject should be alone. This was done for two reasons. 1) This enables more straightforward comparisons to other object-in-context SOR variants, where no conspecific is present. 2) The conspecific partner’s behaviour could bias the experimental subject. As the conspecific partner will have only been present in one out of two of the exposure phases, one of the objects would be unfamiliar for that partner (hence novel) if they were to be present in the test phase. This is crucially different to what the experimental subject has experienced, interacting with both objects during the exposure phases (i.e., both objects should be familiar at test for the experimental subject). Indeed, even though conspecific presence has been previously seen to enhance behavioural expression of learning and memory in rodents^44, 45^, our SOR protocol differs to these where all animals had had the same experience in SOR exposure and or test phases^44, 45^. Moreover, a subordinate’s behaviour could be constrained by a dominant conspecific’s scent-marking or aggression at test^46^, and this in combination with the exposure phase difference could mask the experimental subject’s own strategic preference (or lack thereof).

Mice spent significantly more time exploring the novel object-in-context (context^novel^) configuration (M = 46.97s, SD = 16.77s) than the context^familiar^ configuration (M = 35.14s, SD = 15.62s; *t*_(9)_ = −2.43, *p* = 0.038, *d* = −0.77, CI 95% − 0.04 to −1.46; **Fig. 3A**, left). In contrast, there was no difference between the recency^novel^ (M = 41.02s, SD = 14.13s) and the recency^familiar^ configurations (M = 41.09s, SD 19.93s; *t*_(9)_ = 0.01, *p* =0.99, *d* = 0.004, **Fig. 3A**, right). The discrimination ratio 2 (D2) score data yielded a similar picture (context D2: M= 0.13, SD = 0.22; *t*_(9)_ = 1.83, *p* = 0.10, *d* = 0.58; recency D2: M = −0.005, SD = 0.23; *t*_(9)_ = −0.065, *p* = 0.95, *d* = −0.02). In fact, however, closer inspection showed that when tested in the 1^st^ context, the average context D2 score was positive and strongly different from zero (**Fig. 3B**; M = 0.29, SD = 0.16; *t*_(7)_ = 5.05, *p* = 0.001, *d* = 1.79, CI 95% 0.62 to 2.91), whereas this was not the case for testing in the 2^nd^ context (**Fig. 3C**; M = 0.12, SD = 0.25; *t*_(9)_ = 1.51, *p* = 0.17). One possible explanation for this is relative recency of the contextually specifying cue^28, 43^, which in this case the conspecific was more recently seen in test in the 1^st^ context trials compared to test in the 2^nd^ context trials, potentially making their absence at test more salient in such trials. Yet, this emphasises the importance of context-based SOR research in reporting trial types separately to better understand the possible differences in recognition behaviour between them^27, 28, 43^.

**Figure 3.**
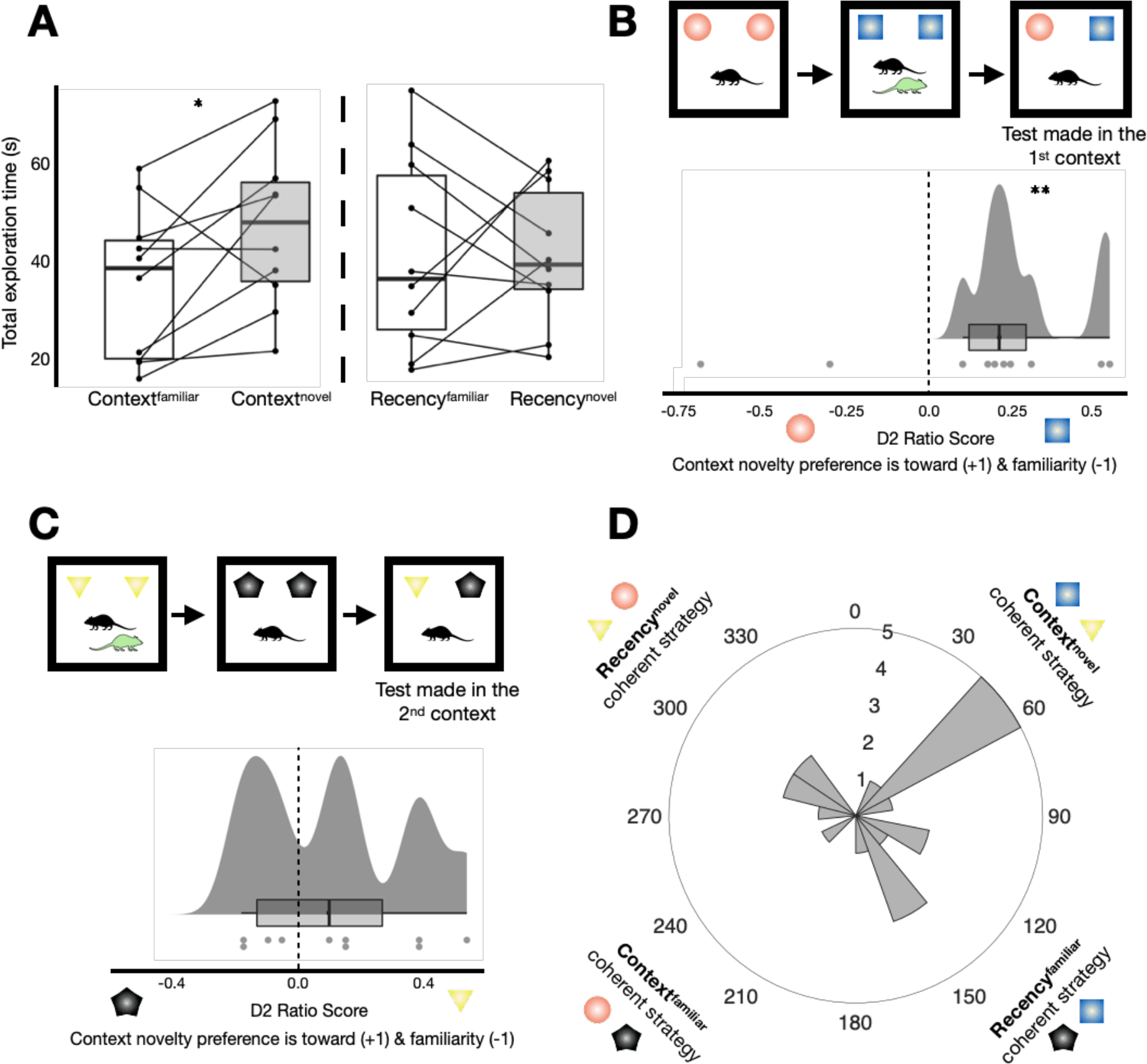
Experiment 1: conspecific presence and absence are sufficient to act as a contextual cue for mice to remember episodes. (**A**) Total exploration time (s) summed across all test phases. Mice explored the context^novel^ configuration (M = 46.97s, SD = 16.77s) significantly more on average than the context^familiar^ configuration (M = 35.14s, SD = 15.62s; *t*(9) = −2.43, *p* = 0.038, *d* = −0.77, CI 95% 0.04 to −1.46). There was no difference between the recency configurations (recency^novel^: M = 41.02, SD = 14.13s; recency^familiar^: 41.09s, SD = 19.93s; *t*(9) = 0.01, *p* = 0.99, *d* = 0.004). (**B**) Overall performance was particularly driven by tests situated in the 1^st^ context. The average context discrimination 2 (D2) score was positive (M = 0.29, SD = 0.16) and differed significantly from zero (*t*(7) = 5.05, *p* = 0.001, *d* = 1.79, CI 95% 0.62 to 2.91). (**C**) The average context D2 score for trials when the test made in the 2^nd^ context was also positive (M = 0.12, SD = 0.25) but did not differ from zero (*t*(9) = 1.51, *p* = 0.17, *d* = 0.48). (**D**) Angular histogram depicting the circular data; n = 20. D2 ratio scores were taken from consecutive trials to form circular data points and thus represents animal-trial data; see methods). Plotted in 16 bins of 22.5°, circular descriptive and inferential statistics are reported in the main text. *Denotes *p* < 0.05. **Denotes *p* = 0.001.

We next asked whether the circular data was uniformly distributed around the circle or whether there was indication of directionality (**Fig. 1**). There was evidence that the circular data was not uniformly distributed around the circle with some biasing towards the context^novel^ quadrant (**Fig 3D**; n = 20, *θ* = 79.9°, *v* = 70.0°, *R =* 0.25, Rao’s spacing test: *U* = 165.31, *p* < 0.05). These results overall suggested that mice used a context-based recognition strategy expressed via novelty preference. This was with performance being mainly driven from test in the 1^st^ context trials, although there was some evidence of coherent context^novel^ object exploration across consecutive trials (that is, also across different trial types; **Fig. 3B-D**). Thus, mice are able to use conspecific presence and their absence as contextual information to separate and identify unique episodes in memory.

### Experiment 2: no coherent strategy emerges when context is specified via an additional local object

This object-in-context task variant used presence and absence of an additional local object as contextual information. Similarly to the dyads of mice, the object acting as a potential context specifier was kept the same throughout the experimental session, as was the physical environment. And mice were only tested in the absence of the object that potentially acted as a context-specifier, in order to be comparable to the conspecific-context variant (experiment 1). Also, this experiment was conducted at the end of the experimental timeline (**SFig. 1**), as this aimed to minimise tedium and possible behavioural carryover affects from the previous object-in-context SOR task^47^.

There was no difference between the total time spent exploring the context^novel^ configuration (**Fig. 4A**, left; M = 42.83s, SD = 21.63s) and the context^familiar^ configuration (M = 62.23s, SD = 29.09s; *t*_(8)_ = 1.47, *p* = 0.18, *d* = 0.49). In addition, there was no difference between the recency^novel^ (**Fig. 4A**, right; M = 61.72s, SD = 39.96s) and the recency^familiar^ configurations (M = 52.70s, SD = 17.87s; *t*_(9)_ = −0.64, *p* = 0.54, *d* = −0.20). The D2 ratio data conveyed a similar picture (context D2: M = −0.03, SD = 0.30; *t*_(9)_ = −0.32, *p* = 0.76, *d* = −0.10; recency D2: M = −0.0003, SD = 0.20; *t*_(9)_ = −0.005, *p* = 1.00, *d* = −0.001). When analysing the different trial types separately, the average context D2 scores for both when the test was situated in the 1^st^ context (**Fig. 4B**; M = −0.03, SD = 0.25), and when situated in the 2^nd^ context (**Fig. 4C**; M = − 0.03, SD = 0.44) were clearly not different from zero (*t*_(9)_ = −0.38, *p* = 0.72, *d* = −0.12; *t*_(9)_ = −0.22, *p* = 0.83, *d* = −0.07; respectively). Lastly, the circular data (**Fig. 4D**; n = 20, *θ* = 172.5°, *v* = 74.2°, *R =* 0.16) was uniformly distributed around the circle (Rao’s spacing test: *U* = 138.84, *p* > 0.50). Therefore, these results suggested that there was no coherent strategy used in the ‘additional local object as context’ variant.

**Figure 4.**
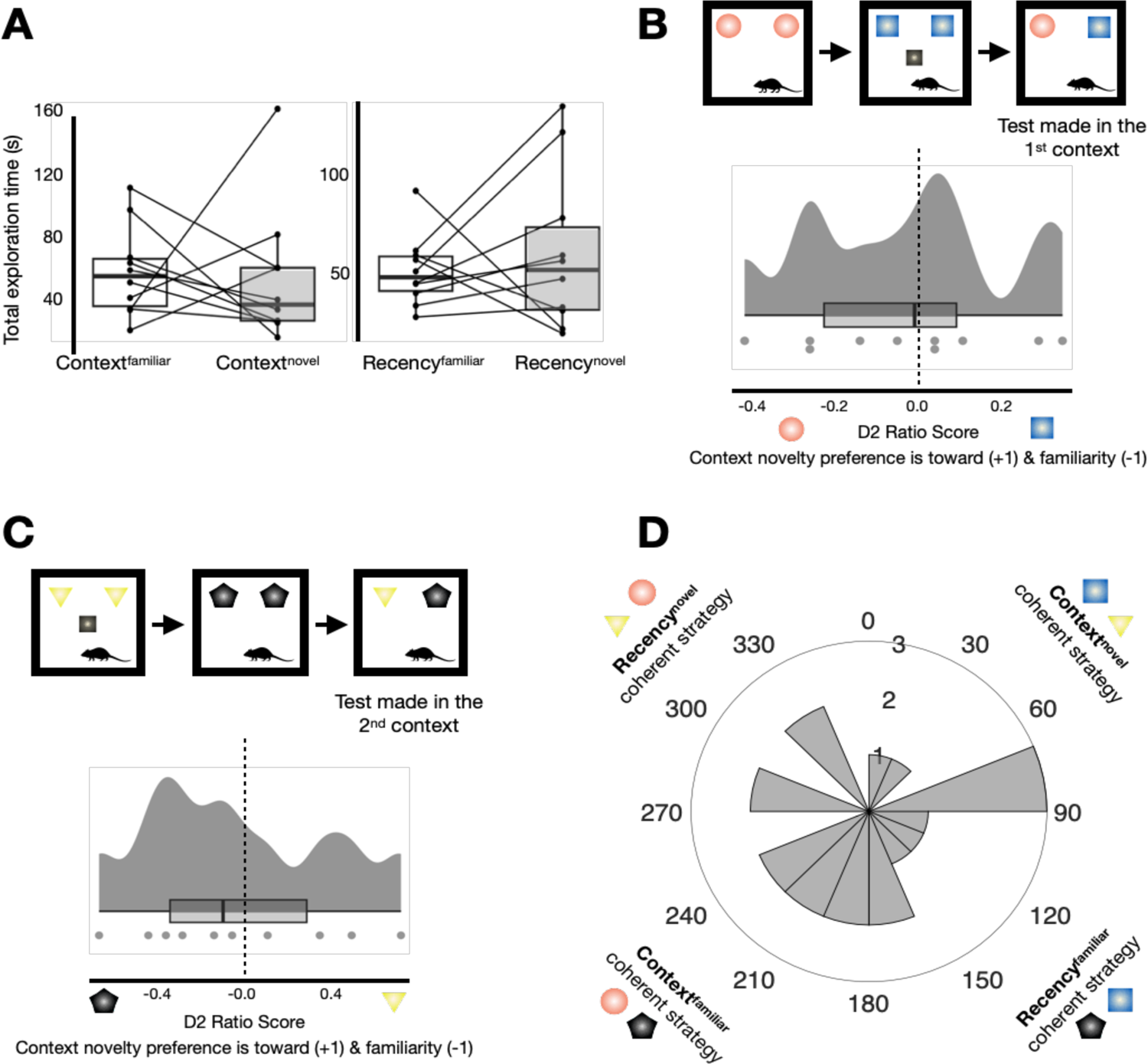
Experiment 2: no coherent strategy emerges when context is specified via an additional local object. (**A**) Total exploration time (s) summed across all test phases. There were no differences between exploration of any particular configurations (context^novel^: M = 42.83s, SD = 21.63s; context^familiar^: M = 62.23s, SD = 29.09s; *t*(8) = 1.47, *p* = 0.18, *d* = 0.49; recency^novel^: M = 61.72s, SD = 39.96s; recency^familiar^: M = 52.70s, SD = 17.87s; *t*(9) = −0.64, p = 0.54, *d* = −0.20). (**B**) The average test in the 1^st^ context D2 score did not significantly from zero (M = −0.03, SD = 0.25; *t*(9) = −0.38, *p* = 0.72, *d* = −0.12). (**C**) The average test in the 2^nd^ context D2 score did not significantly from zero (M = −0.03, SD = 0.44; *t*(9) = −0.22, *p* = 0.83, *d =* −0.07). (**D**) Angular histogram depicting the circular data (n = 20), plotted in 16 bins of 22.5°, circular descriptive and inferential statistics are reported in the main text. Of note, the objects used in this task variant were not the same as used in the conspecific as context task variant (i.e., the schematics are kept the same for clarity purposes).

### Comparison of the context specifiers: conspecific partner (experiment 1) and the additional local object (experiment 2)

Due to our protocol of testing the subject when they were only alone, another possibility that could explain recognition behaviour (during test phases of these object-in-context variants), is a simple object recognition strategy based upon novelty-detection with ignorance of the contextual information. For example, this could occur if there was little acquisition of objects when exposed in the presence of the conspecific relative to when the subject was alone. Hence, mice would explore the same object as predicted via a context^novel^ strategy but due to this object simply being more unfamiliar (and thus novel) at test. We thus conducted control analyses concerning the object exploration in exposure phases of experiment 1 and experiment 2 (**SFig. 2**).

A mixed repeated measures ANOVA comparing summed total exploration in exposure phases relative to test phases, revealed that for both experiment 1 and experiment 2 there was more exploration in exposure phases vs. test phases (**SFig. 2A-B**; exploration of exposure phases was scaled to match that of test phases; Fisher’s least significant difference, LSD, post-hoc tests: *p* = 0.01, *p* < 0.001, experiment 1 and 2; respectively). This suggested successful acquisition of objects did occur during exposure phases; in other words, objects at the test phases were likely familiar to mice in both experiments 1 and 2. Interestingly, we also found significantly more total exploration on average in experiment 2 (M = 143.71) relative to experiment 1 (M = 101.72; F_(1,9)_ = 14.04, *p* = 0.005, η_p_^2^ = 0.61), indicating that there was minimal decline in task motivation across experiments 1 and 2.

We next sought to compare exposure phases of the ‘context’ specifiers (i.e., conspecific partners in experiment 1 vs. an additional local object in experiment 2), and ‘presence’ of the context specifier (that is, the context specifier’s presence vs. its absence). A mixed repeated measures ANOVA yielded a significant interaction between ‘context’ and ‘presence’ (F_(1,9)_ = 9.11, *p* = 0.02, η_p_^2^ = 0.50). Fisher’s LSD post-hoc analyses indicated that within experiment 1, there was significantly more object exploration in exposure phases when the conspecific was present (**SFig. 2C**; M = 34.59) versus when mice were alone (M = 26.08; *p* = 0.017), which is in accordance with previous reports^44, 45^. Contrastingly, within experiment 2, levels of exploration in the presence of the additional local object (M = 40.69) were similar to when it was absent (M = 45.82; *p* = 0.32).

In summary, the control analyses further support the notion that in experiment 1 mice were using a mnemonic strategy reliant on the contextual information (the conspecific partner; **Fig 3**). However, in experiment 2, despite some indication of successful object acquisition from exposure phases (similarly to that seen in experiment 1; **SFig. 2A-B**) we found no evidence of a coherent recency-based or context-based strategy when an additional local object could have been used as a potential context specifier (**Fig. 4**).

### Experiment 3: mice preferentially explore contextual mismatch information associated with familiar conspecifics over a recency-based mnemonic strategy

Inspired by the object-in-context SOR paradigm, for experiment 3, we adapted the standard social discrimination task^22, 23^ to construct a conspecific-in-context variant (see Introduction, **Fig. 2, SFig. 3**). The aim of the design was to make two novelty-oriented discriminatory strategies available, and to pit them against each other. Figure 2 pictorially illustrates the two potential strategies. In the test phase, the mice could preferentially explore either: 1) the conspecific in the novel conspecific-in-context configuration, seen more recently and presented in the same place, but where there was now a contextual mismatch (conspecific C, yellow, in **Fig. 2**); or 2) the conspecific who was seen longer ago, presented in the same place and context (conspecific A, red, in **Fig. 2**, recency strategy). In this way, we could investigate the question of whether mice show a spontaneous exploratory preference for a context-based or recency-based mnemonic strategy when both are available, as in the context SOR tasks but now with respect to conspecifics.

Experiment 3 comprised three sessions of this conspecific-in-context design, with a single trial per session. Two sessions were with conspecifics of the same sex, and one of the opposite-sex (**SFig. 1**). Having a second same-sex session allowed for examination of recognition behaviour once subjects had had further habituation of the task conditions, whilst also allowing for within-subject counterbalancing of context order and conspecific placement to enhance within-subject reliability.

Moreover, previous work in rodents has suggested that social interaction behaviour and neuromodulatory mechanisms can be dependent on conspecific-sex, with increased salience associated with members of the opposite sex^48–50^. Thus, the idea of the final opposite-sex session was to examine whether recognition behaviour would differ because of using opposite-sex conspecifics which should be more socially salient stimuli. In this way, the to-be-recognised opposite-sex conspecifics may hinder or boost preferential exploratory behaviour (of a particular recognition strategy) relative to same-sex conspecific stimuli.

A mixed repeated measures ANOVA was conducted on the exploration behaviour in the test phases across sessions (**Fig. 5A**). There was an overall significant main effect of ‘session’ (same-sex sessions 1 and 2, and the opposite-sex session 3; F_(1.24,11.13)_ = 26.73, *p* < 0.001, η_p_^2^ = 0.75). Bonferroni corrected post-hoc tests showed that there were comparable levels of exploration across same-sex session 1 (M = 4.97s) and same-sex session 2 (M = 4.13s, *p* = 1.00). Whereas there was significantly more exploration in the opposite-sex session 3 (M = 18.08s) relative to session 1 and 2 (*p* = 0.003, *p* < 0.001; respectively). There was also an overall significant main effect of ‘conspecific’ (that is, conspecific A, red, vs. conspecific C, yellow, see **Fig. 2** and **5**, F_(1,9)_ = 5.67, *p* = 0.04, η_p_^2^ = 0.39). Post-hoc tests showed that across sessions there was more exploration of the contextually mismatched conspecific C (M = 10.34s; **Fig. 5B**) relative to the least recently seen conspecific A (M = 7.78s, *p* = 0.04) who was presented in the same context and place at test. This is consistent with forming an episodic-like conspecific-in-context memory.

**Figure 5.**
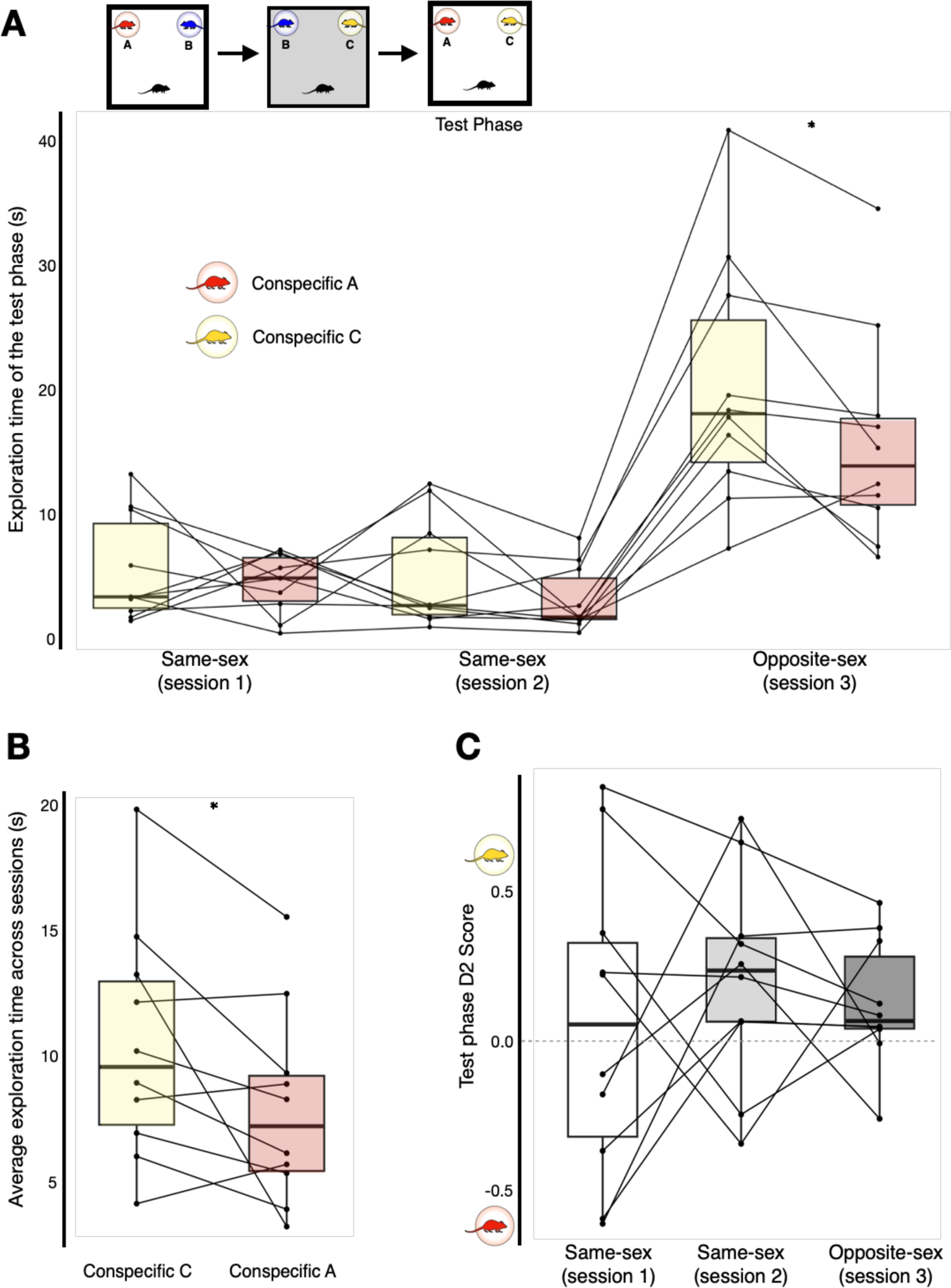
Experiment 3: mice preferentially explore contextual mismatch information associated with familiar conspecifics over a recency-based mnemonic strategy. (**A**) Upper left: reminder schematic of the conspecific-in-context task (see also Fig. 2). Lower: exploration times of conspecific A and C in the test phase by session. (**B**) Average exploration time of conspecific A (M = 7.78s) and C (M = 10.34s) across all sessions. (**C**) D2 ratio score of the test phase by session. (A-C) Descriptive and inferential statistics reported in the main text. *Denotes *p* <0.05.

There was no overall interaction between session and conspecific (F_(2,18)_ = 1.32, *p* = 0.29, η_p_^2^ = 0.13). Yet, similarly to the overall significant main effect of session, post-hoc tests showed that regardless of conspecific (A or C) more exploration was made session in 3 relative to session 1 and 2 (all *p* ≤ 0.006; comparable exploration levels across session 1 and 2, all *p* ≥ 0.60. This very clear result suggests the enhanced salience of members of the opposite-sex. Given these marked differences in exploratory expenditure across same-sex sessions versus the opposite-sex session, we next sought to check using the D2 ratio score (**Fig. 5C**; which accounts for individual differences in exploration levels), whether preferential exploration towards certain conspecifics differed across sessions. A repeated measures ANOVA yielded no sign at all of differences in recognition performance across sessions (F_(2, 18),_ *p* = 0.64, η_p_^2^ = 0.05, Bonferroni post-hoc tests all *p* = 1.00).

This suggested that exploratory preference on average was similar across these sessions, validating our finding of overall exploratory preference towards conspecific C (**Fig. 5B**). Finally, post-hoc tests within sessions, revealed that there was comparable levels of exploration towards conspecific A and C in session 1 (Conspecific C: M = 5.52s, Conspecific A: M = 4.42s, *p* = 0.53; D2 score: M = 0.06, SD = 0.52, *t*_(9)_ = 0.34, *p* = 0.75, *d* = 0.11), and in session 2 (Conspecific C: M = 5.19s, Conspecific A: M = 3.08s, *p* = 0.12; D2 score: M = 0.21, SD = 0.35, *t*_(9)_ = 1.89, *p* = 0.09, *d* = 0.60, CI 95% −0.09 to 1.26). However, in the opposite-sex session, there was significantly more exploration of conspecific C (M = 20.32) versus conspecific A (M = 15.84, *p* = 0.044; D2 score: M = 0.12, SD = 0.21, *t*_(9)_ = 1.82, *p* = 0.10, *d* = 0.58, CI 95% −0.11 to 1.24). This suggested that the overall exploratory preference toward the contextually-mismatched conspecific C (**Fig. 5B**) was particularly driven by recognition behaviour in the opposite-sex session, possibly due to the enhanced salience in the nature of the social stimuli^48–50^.

## Discussion

Being able to flexibly remember episodes via social-context is of evolutionary importance^1-4^. Our experiments suggest that the same cohort of mice not only preferentially explored conspecifics associated with environment-based contextual mismatch information (experiment 3; **Fig. 5**), but used conspecific presence and absence as a means to remember unique episodes in memory (experiment 1; **Fig. 3**). These findings echo the substantial evidence reported using SOR paradigms, that when there is availability of both context-based novelty and recency-based novelty, rodent exploratory behaviour is more directed to the unexpected contextual change^28–30^.

Interestingly, no overall strategy emerged when context was specified by an additional local object (experiment 2; **Fig. 4**), and this was seemingly not due to reduced motivation nor lack of object acquisition during exposure phases (**SFig. 2**). There are undeniable differences between conspecifics and a static local object, in terms of the sensory cues that emanate from them and their biological relavance^1,48–50^. Yet, it would of course be premature to conclude from this experiment alone that objects cannot be used by mice as contextual information in defining unique episodes (especially when learnt via explicit reinforcing^14, 31^). What does seem reasonable to conclude is that, the presence/absence of a conspecific has sufficient ethological salience under incidental spontaneous conditions to be incorporated into episodic-like memory, and this salience is clearly greater than that for a man-made object, (especially when that object becomes increasingly habituated to over time, as was the case in experiment 2).

A previous study suggested that when rats were exposed to an unfamiliar context (a change in the physical environment), there was a reduction of investigation and mild aggression towards a juvenile conspecific, who was increasingly familiarised to from three previous sessions in a different, familiarised context^51^. The experimenters argued that although rats still recognised the conspecific, the behavioural change could be interpreted as increased habituation to the conspecific^51^, that is, not only due to contextual novelty but perhaps a novel association of the conspecific-in-context. We extend such work by showing in experiment 3, that when mice are given a free choice to explore a more recently-seen familiar conspecific associated with environment-based contextual mismatch and a less recently seen familiar conspecific presented in the same place and context, they preferentially explore the former.

Converging evidence demonstrates that successful social mnemonic processing can strongly rely upon the hippocampal formation^52–56^. Hippocampal principal cells can show place-dependent activity as rodents traverse their environment, hence termed place cells^33–35^. But strikingly, hippocampal principal cells may also flexibly integrate information about conspecifics in their responsivity^56–61^, for example place-like activity of these cells can also relate to positional information of conspecifics (i.e., social place cells^58–60^). Notably, such social place cells were not reported when rats’ behaviour was dependent upon observationally tracking a robot’s movement^62^.

When substantial changes are made to the environment, place cells exhibit a phenomenon known as ‘remapping’, whereby some cells fire in one environment, but not another, or fire in different locations in each environment^33–36, 63–65^. Thus, at the population-level, two sufficiently-different environments are represented distinctly, via ‘global remapping’^35, 63^, in a manner that may specify two different contexts. Indeed, the argument has been explicitly made, potentially finessing long-running issues with defining ‘context’, that “electrophysiology opens the door to a measurement-based approach with a clear definition: a new context is one that is sufficient to evoke global remapping”^66^.

It seems reasonable to infer that hippocampal place cell remapping occurs in the majority of context-SOR studies, since these studies typically employ marked changes in the physical environment to specify context. Moreover, increases in rearing on hind legs typically accompanies place cell remapping in novel, physically different, contexts^67–69^ implying a link between place cell remapping and context-sensitive exploratory behaviour in rodents. How strong place cell remapping needs to be, and in which hippocampal sub-regions, to act as a universal context-differentiation readout signal remains unclear. Our behavioural observations here suggest that conspecific presence/absence can define ‘context’ and thus distinguish otherwise-similar episodes in the same physical environment. Taken together with evidence of partial place cell remapping in sub-regions CA2 and ventral CA1 across scenarios that differ only socially^54, 70, 71^ it seems reasonable to suggest that: 1) social as well as physical-environmental cues can define behaviourally-relevant context shifts in rodents as well as other species, especially humans; 2) remapping in hippocampal place cells, even in rodents, may not need to be driven by changes in physical-environmental cues, nor to be ‘complete/global’, in order to serve as a context-shift signal.

In conclusion, we have implemented new spontaneous recognition task variants to show that mice readily use social episodic-like memory to drive their exploration. The tasks offer a novel way to tease apart the mechanisms of social recognition memory in a crucially different way to the current, frequently used social discrimination protocols. This is especially relevant in modelling atypical, disease and neuropsychological disorders with rodent models, as in response to given manipulations animals may display chance level performance or be using recency-based or context-based (episodic-like) mnemonic strategies to guide their behaviour.

## Methods

### Subjects

Ten B6FVBF1 mice (5 male) were bred inhouse at the life science support unit (Durham University, U.K.). They were ∼10 weeks of age when habituation begun (Females weight: M = 24.1g, SD = 1.0g; Males weight: M = 30.4g, SD = 1.8g) and were housed in two cages in same-sex groups of 5. Each home cage measured 45 × 28 × 13 cm (l × w × h; Model: MB1, NKP isotec., U.K.) and were equipped with 2 mouse tunnels and 2 igloos (Datesand Limited., U.K.). The home room was maintained on a 12-h light-dark cycle (07:00-19:00h), with daily monitoring of temperature and humidity (20 ± 1°C; 55 ± 10%; respectively). All stages occurred during the light phase and mice had free availability of food and water ad libitum throughout (i.e., were not food or water deprived). Animals were not euthanised as part of the experiments. All experiments were conducted in accordance with the U.K. Animals Scientific Procedures Act (1986), approved by Durham University AWERB and in accordance with the Home Office (procedure licence number: P7B7D2E4B). Reporting follows the recommendations in the ARRIVE guidelines.

### Apparatus and objects

All reported experiments took place in an apparatus designed for spontaneous recognition (Model CI.80514R-1, Campden Instruments., U.K.). The specifications of which are previously reported^27^. Only the open field area was used presently, white noise played continuously from above the open area (62 ± 8.5 dB SPL) and an additional camera was also used for behavioural recording (Model: MWC72ZD/A, iPhone 11 Pro). Environmental contexts were comprised of sensorily distinct floors and were sometimes paired with a wall cue (see SFig. 3A-C). The objects varied in material, shape, size, texture and visual complexion, each object had a minimum of 3 duplicates and were paired quasi-randomly (example pair shown in SFig. 3D). In experiment 2, the additional local object that could act as context is shown in SFig. 3E. For experiment 3 the social conspecific-in-context recognition experiments, conspecifics were placed within a wire cup, and all were weighed down with the same object (see SFig. 3F). For all experiments, objects and conspecifics were positioned towards the far corners of the open field opposite the door (i.e., mice egocentrically had objects/conspecifics left and right to them, as they were placed into the open field, always in the same direction, north towards the objects/conspecifics). At the end of testing sessions for the experiments 1 and 2, objects, floors and the apparatus were cleaned using disinfectant wipes (Clinell universal wipes, GAMA Healthcare Ltd., U.K.). For experiment 3, the wire cups and floors were cleaned and dried between each phase and at the end, this was to minimise the crossing of scent-marking cues of conspecifics between phases.

### Habituation

Mice were first handled in their home room for a minimum of 3 consecutive days, before being transported (in cage groups) to the experimental room where all reported testing took place (white noise played and the room was lit by diffuse white light from 2 lamps, 60 W & 100 W). The first-time mice were habituated to the open field was in context X (see SFig. 3A) and they did so in cage groups (30 minutes). Following this, they were habituated once in the same dyads as used for the experimental session, but objects were now present (two of the same and they were not used in any experiments; 30 minutes). Prior to experiment 3, context Y and Z (SFig. 3B-C) were habituated to on the same day in cage groups (30 minutes each, ∼1.5 hours between; the wire cups were present). Lastly, prior to experiment 2, context X was re-habituated twice on separate days, once without objects and once with the same habituation objects as used previously. Both habituations occurred in cage groups and lasted for 20 minutes.

### Procedure: object-in-context experiments (experiments 1 and 2)

A given trial was composed of 3 phases (Fig 1A; 2 exposure phases and a test phase). The same pair of objects are placed in exposure 1, where mice explored them for ∼3 minutes before being returned to a separate holding cage (for ∼3 minutes, the same design as the home cage and kept within the experimental room). A different pair of objects are presented in exposure 2 and again mice explored them for ∼3 minutes. Approximately 5 minutes elapsed before experiencing of the test phase, which contained a copy of an object from exposure 1 and a copy of an object from exposure 2 and lasted for ∼3 minutes (example object pair shown in SFig. 3D). There was a ∼3 minute interval before the next trial begun. For experiment 1, a given dyad of mice were composed randomly of same-sex cage (and litter) mates. The test subject was always placed into the context first and removed last, being returned into the same holding cage as the partner. The partner who was not the test-subject for that session was tested 6 days later (SFig. 1). For experiment 2, the additional local object acting as a potential context specifier (see SFig. 3E) was placed in with the other objects but kept the same throughout the session and was never present in the test phase (like in experiment 1). The objects used in experiment 2 were not the same as used in experiment 1. Test phases could be situated in the 1^st^ context or test phases could be situated in the 2^nd^ context (Fig. 1A). Four trials comprised a single experimental session and the trial order, object order and placement of the object-in-context novelty was counterbalanced. Notably, the context specifier could not be counterbalanced as we required the subject to always be alone in the test phase (see main text for reasoning).

### Procedure: social conspecific-in-context recognition (experiment 3)

A given trial was composed of 3 phases (2 exposure phases and a test phase; see Fig. 2). In the first exposure phase, the test subject experienced two conspecifics contained within wire cups in a given environment-based context (∼3 minutes), before being returned alone into a holding cage. After ∼5 minutes, the test subject was placed back into the apparatus but now the context had been changed and they could explore a familiar conspecific or a newly introduced unfamiliar conspecific in the task conditions altogether (∼3 minutes). This was considered as a second exposure phase to allow for the scenario that occurs in the test phase. Again after ∼5 minutes elapsed, the test subject was returned into the apparatus and the context was changed back to that experienced in the first exposure phase. Subjects in the test phase could then explore a familiar conspecific, more recently-seen in the same place, who now had a contextual mismatch or they could explore an also familiar conspecific who was seen less recently, but presented in the same place and context (lasting ∼3 minutes). This experiment was conducted twice using same-sex cage and littermates (the second session was within-subject counterbalanced) and lastly once using opposite-sex littermates (SFig. 1; all randomly assigned as to which conspecifics were to-be-recognised, and all mice experienced containing in the wire cups, within completion of sessions across animals). The context and conspecific order were all counterbalanced and hence so was the placement of the novel conspecific-in-context in test trials.

### Behavioural analyses

Behaviour was measured off-line via the recorded footage of experimental trials. Exploratory behaviour was regarded as when mice were within ∼2cm of the object (or the wire cup/conspecific) and actively exploring it (i.e., sniffing, touching, biting and visibly whisking). Behaviour such as climbing and sitting upon objects, or the wire cup configurations were not considered as exploration, and neither was using them to support rearing. The duration of exploratory behaviour (s) with respect to objects, wire cups and conspecifics (of all phases) was manually scored unblinded by the main experimenter (#1). All reported statistics are based upon the main experimenter’s scoring. Importantly, a random subset (20% of each experiment test phase) was scored blinded by two other trained experimenters (#2 and #3, who had less experience overall in comparison to experimenter #1). Scoring between all experimenters were significantly and positively correlated (#1 vs. #2: *r*(54) = 0.75, *p* < 0.001, CI 95% 0.60 to 0.85; #1 vs. #3: *r*(54) = 0.83, *p* < 0.001, CI 95% 0.72 to 0.90; #2 vs. #3: *r*(54) = 0.89, *p* < 0.001, CI 95% 0.81 to 0.93). Additionally, intraclass correlation coefficient (ICC) analysis suggested good to excellent reliability of scoring^72^ (The average measure ICC was 0.91, CI 95% 0.77 to 0.96, F(55,110) = 19.07, *p* < 0.001; 2-way random-effects model, absolute-agreement, *k* = 3, mean-rating).

Total exploration time (s) was the summed exploration across trials by animals of a given configuration (e.g., the novel object-in-context) or else specified. In no case did side-bias better explain recognition performance over a context or recency-based strategy (Fig. 3A; Fig. 4A; Fig. 5C; *t*(9) = −1.92, *p* = 0.09; *t*(8) = −0.12, *p* = 0.90; *t*(9) = 0.86, *p* = 0.42; respectively). The classically described discrimination ratio 2 (D2) scores for a context-based or recency-based strategy is previously reported^27^, from a context D2 ratio score calculation novelty preference is toward +1 and familiarity preference toward −1, with 0 indicating no preference. For experiment 3 D2 scores, preference to explore the conspecific associated with the contextual mismatch was indicated as values towards +1. For each animal, the D2 score was calculated individually for each trial and then averaged across all trials and finally across animals to give the reported overall mean D2 scores (unless specified by trial/test type). All data was tested for normality (& sphericity where applicable) and a non-parametric alternative (Greenhouse-Geisser correction) was used if *p* < 0.05, using SPSS, v28 (2021, IBM Corp). Outlier cases were identified based on quartiles (where *k* = 2.07)^73^ and were excluded from statistical tests. All reported measures were two-tailed tests.

We plotted animals’ test in the 1^st^ context D2 scores against test in the 2^nd^ context D2 scores and formulated circular data (via an arctangent function, converted from radians to degrees, 0 ± 180°). Importantly, such circular analyses should be interpreted with dependence upon evidence of exploratory preference differing to chance level performance, but it can enhance explanatory power of the spontaneous recognition task data. Perfectly coherent strategies across trial types (Fig. 1B) would be indicated by circular data points aligning at 45°, 135°, −135° (225°) and −45° (315°). For example, ∼45° for a coherent context^novel^ strategy in a context-based SOR task (Fig. 1C). Data points aligning more towards 0°, 90°, 180° and −90° (−270°) would suggest that exploratory preference is exhibited in only 1 out of the 2 trial/test types. We designed the object-in-context experiments (experiments 1 and 2) in such a way where test in the 1^st^ context trials were interleaved with test in the 2^nd^ context trials, allowing 2 consecutive trials (i.e., the first and last 2 trials from experiment 1 and 2)) to form animal-trial circular data points. We used the MATLAB (2020b, The MathWorks, Inc) circular statistics toolbox^74^ and package circular^75^ in R (2021.09.0, RStudio, PCB) to compute circular descriptive and inferential statistics. To use a single circular test capable of accommodating distributions that were not expected to be unimodal, we employed Rao’s spacing test^74^.

## Data availability

The data that support the findings of this manuscript are available on the OSF repository (https://doi.org/10.17605/OSF.IO/QWZAM) or on available request from T.W.R., the corresponding author.

## Competing interests

The authors declare no competing interests.

## Contributions

All authors contributed to the conceptualisation of the experiments. T.W.R. designed the experiments, collected the data and conducted the data analyses. S.L.P., C.L., and A.E. supervised these stages. All authors interpreted the data. T.W.R. wrote the original manuscript and edited it, and S.L.P., C.L., and A.E. reviewed the original manuscript and edited it.

## Acknowledgements

This work was supported by a BBSRC grant to C.L. (PI) and S.L.P. (co-I) (BB/T014768/1). The authors would also like to thank A. Kitching. and the staff at the life science support unit (Durham, U.K.), and thank B.J.A. Slater. and G. Kan. for their assistance with scoring.

---

**Supplementary Figure 1.**
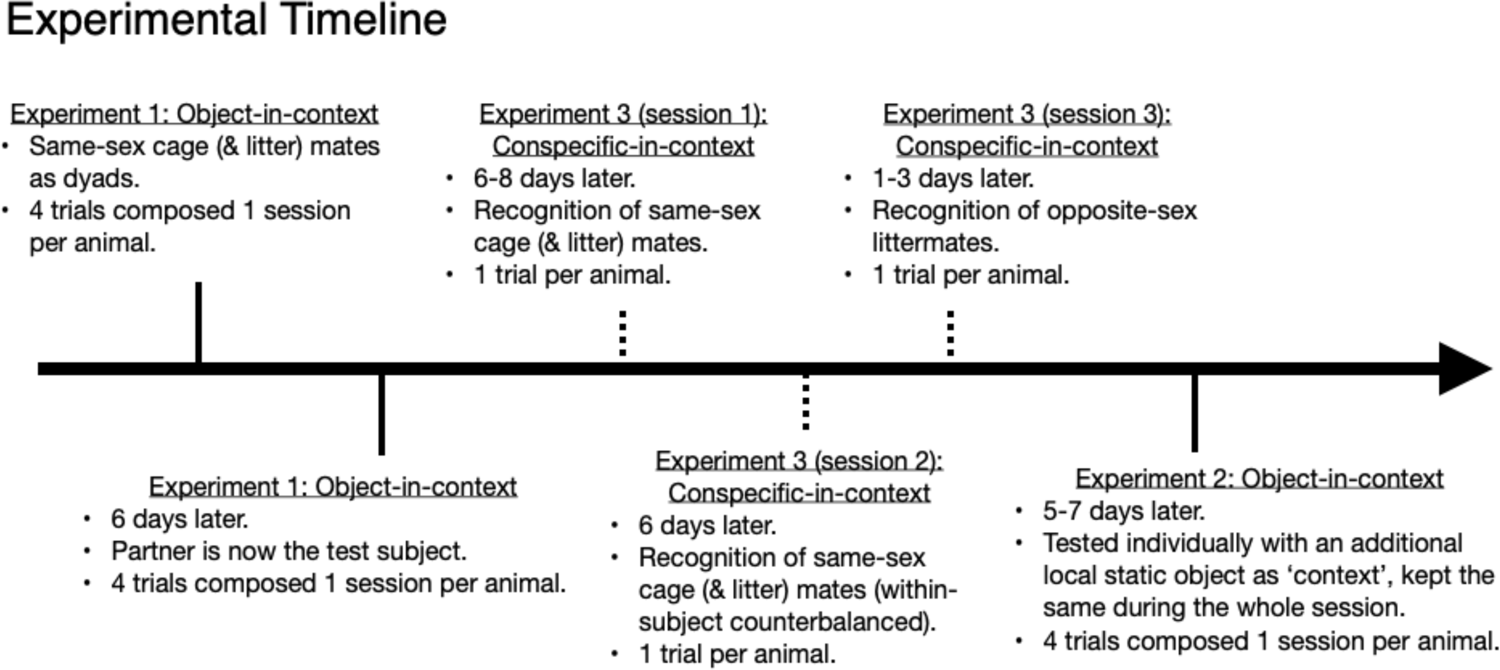
The timeline for all the presently described experiments.

**Supplementary Figure 2.**
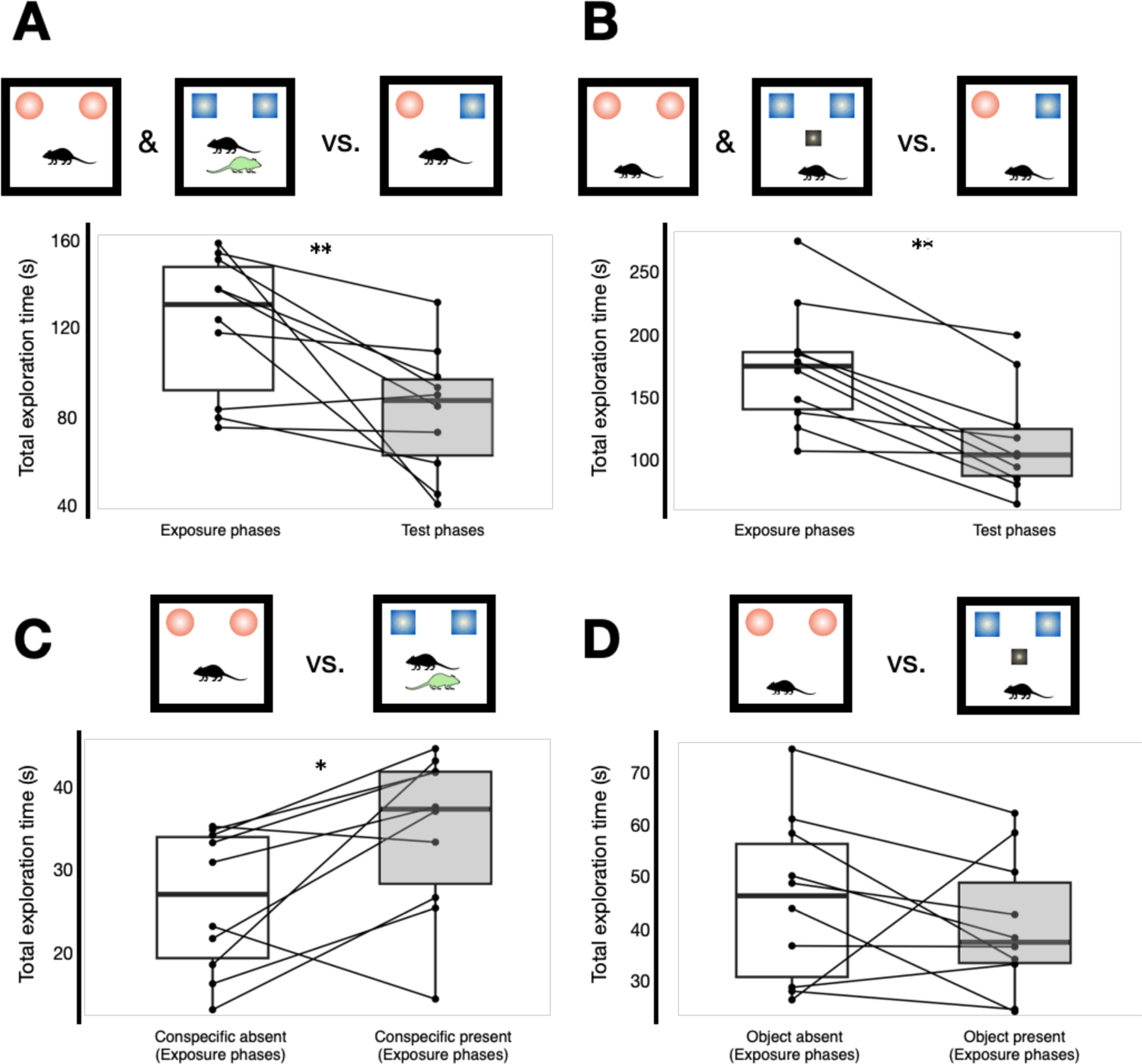
Analyses of object exploration during the exposure phases of the object-in-context variants. (**A**) Summed total object exploration during exposure phases scaled to compare against summed total exploration in the test phases. A mixed repeated measures ANOVA yielded a significant main effect of ‘context’ (conspecific, experiment 1, vs. object variant, experiment 2; F(1,9) = 14.04, *p* = 0.005, ηp^2^ = 0.61). Fisher’s least significant difference post-hoc (LSDph) analyses (for all following comparisons) revealed that there was significantly more exploration in Exp 2 (M = 143.71) vs. Exp 1 (M = 101.72, *p* = 0.005). There was also a significant main effect of ‘phase’ (exposure vs. test phase; F(1,9) = 46.70, *p* < 0.001, ηp^2^ = 0.84. Significantly more exploration in exposure phases (M = 147.17) vs. test phases (M = 98.26, *p* < 0.001). The ANOVA yielded no initial overall interaction between context and phase (F(1,9) = 1.25, *p* = 0.29, ηp^2^ = 0.12). However, post-hoc tests revealed that within context, there was significantly more exploration in the exposure phases of Exp 1 (M = 121.33) vs. the test phases (M = 82.11; *p* = 0.01, *shown in A*). (**B**) Post-hoc tests also revealed that there was significantly more exploration in the exposure phases of Exp 2 (M = 173.01) vs. the test phases (M = 114.42; *p* < 0.001). (**C**) A mixed repeated measures ANOVA was conducted for object exploration during only the exposure phases. Similarly to A, there was a significant main effect of context (F(1,9) = 12.33, *p* = 0.007, ηp^2^ = 0.58; more exploration in Exp 2, M = 43,25, vs. Exp 1, M = 30.33, *p* = 0.007). There was no main effect of ‘presence’ (conspecific/object presence vs. absence; F(1,9) = 0.25, *p* = 0.63, ηp^2^ = 0.03), nor ‘trial-type’ (test made in the 1^st^ context vs. test made in the 2^nd^ context trials; F(1,9) = 1.58, *p* = 0.24, ηp^2^ = 0.15). There was a significant 2-way interaction between context and presence (F(1,9) = 9.11, *p* = 0.02, ηp^2^ = 0.50). Post-hoc tests revealed that within presence, there was more exploration in Exp 2 in absence of the object (M = 45.82) vs. when mice were alone in Exp 1 (M = 26.08, *p* = 0.001). However, there was no difference between the presence exposure phases across Exp 1 and 2 (Exp 2: M = 40.96; Exp 1: M = 34.59, *p* = 0.19). *As shown in C*, within context (of Exp 1), there was significantly more exploration when there was conspecific presence (M = 34.59) vs. their absence (M = 26.08, *p* = 0.017). Finally, the ANOVA revealed no overall 3-way interaction between, context, presence and trial-type (F(1,9) = 0.11, *p* = 0.75, ηp^2^ = 0.01; within context & trial-type: test in the 1^st^ context conspecific presence, M = 32.66 vs. alone, M = 24.64, *p* = 0.09. Test in the 2^nd^ context conspecific presence, M = 36.52 vs. alone, M = 27.52, *p* = 0.11). (**D**) There was no difference between the exposure phases of Exp 2 (presence: M = 40.69 vs. absence: M = 45.82; *p* = 0.32). Within context and trial-type: test in the 1^st^ context trials object acting as context present (M = 44.75), vs. absent (M = 53.53, *p* = 0.43). Test in the 2^nd^ context trials object present (M = 36.62), vs. absent (M = 38.11, *p* = 0.87). Of note, schematics of only test in 1^st^ context trials are shown for consistency, both trial types were considered for all the above reported analyses. *Denotes *p* < 0.05, **Denotes *p* ≤ 0.01

**Supplementary Figure 3.**
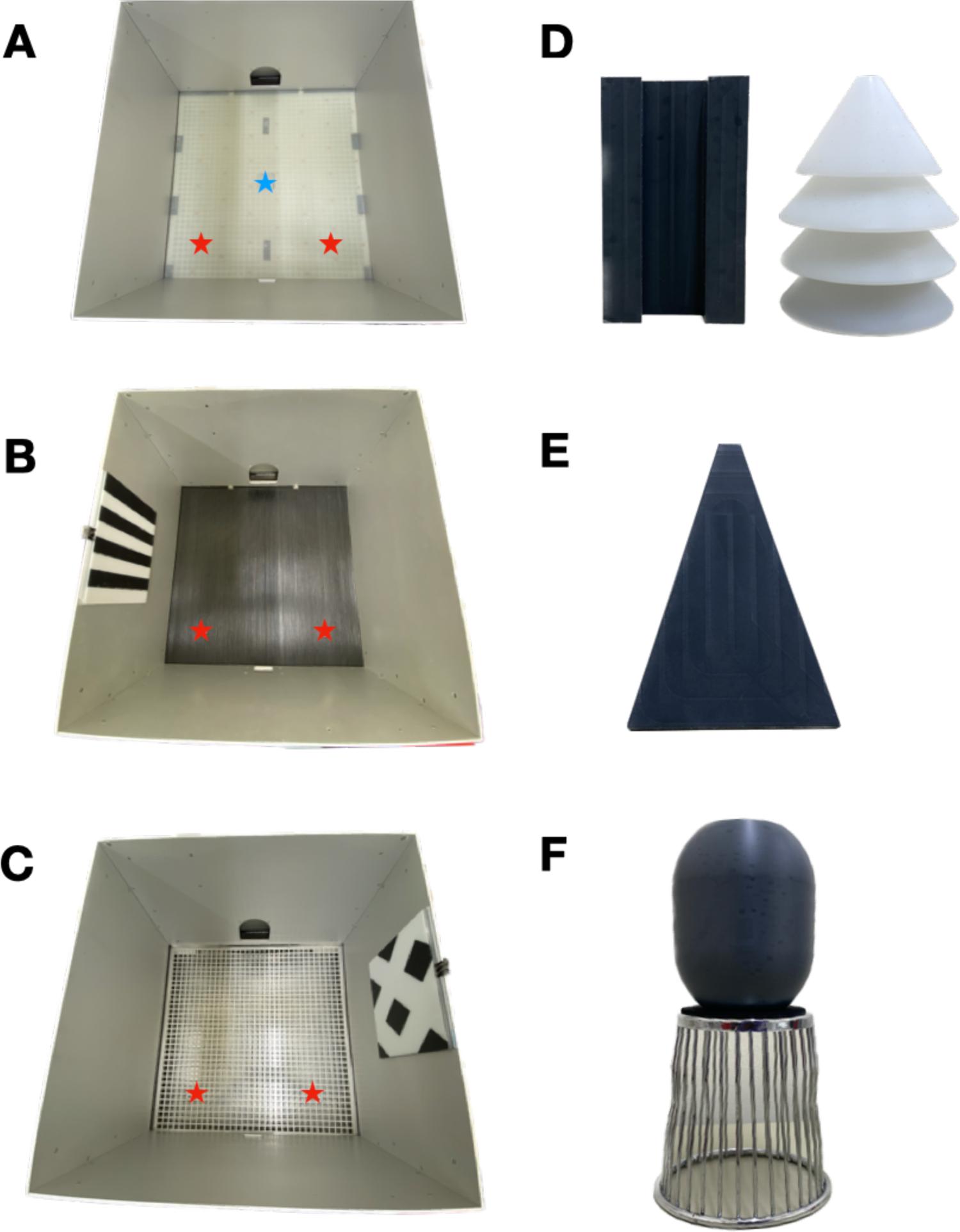
Environment-based contexts and objects. (**A**) Context X open field, used for the object-in-context spontaneous recognition variants (experiment 1 and 2). It was comprised of no wall cues and a translucent Perspex floor with no holes. For reference, the door was considered south and the objects were placed towards the far corners north indicated via the red stars. The blue star indicates placement of the additional local object (see E) that could act as context. (**B**) Context Y open field, one of the two contexts used for the social conspecific-in-context recognition experiment 3. It was comprised of a striped, textured rubber black floor, paired with a polarised striped cue card on the east wall. Red stars indicated approximate placement of the wire cups (see F) containing conspecifics. (**C**) Context Z open field, the other context used for the conspecific-in-context social recognition experiment 3. It was comprised of steel mesh flooring paired with a polarised diamond patterned cue card on the west wall. (**D**) Example object pair used for the object-in-context experiments. Black object: 5.5 × 5.5 × 9.0cm (l × w × h). White object: 8.0cm diameter, 9.0cm height. (**E**) The additional local object acting as context, kept the same throughout the session. Position indicated via the blue star in A. It measured 5.5 × 5.5 × 7.2cm. (**F**) The chrome steel wire cup (10.2cm diameter, 10.8cm height; Model: 31570, Spectrum Diversified Designs, Inc., Ohio, U.S.A.) used to contain conspecifics, and object used to weigh it down (8.0cm diameter, 9.0cm height). See red stars in B and C for approximate placement in the environment. Of note, the lighting during experimental testing was dimmer than that depicted in A-C.

## Notes

https://doi.org/10.17605/OSF.IO/QWZAM

### Competing Interest Statement

The authors have declared no competing interest.

