## Supplementary figures for "Mice remember experiences via conspecific-context: models of social episodic-like memory"

Supplementary figure 1. (p. 2).
Supplementary figure 2. (p. 3).
Supplementary figure 3. (p. 4).

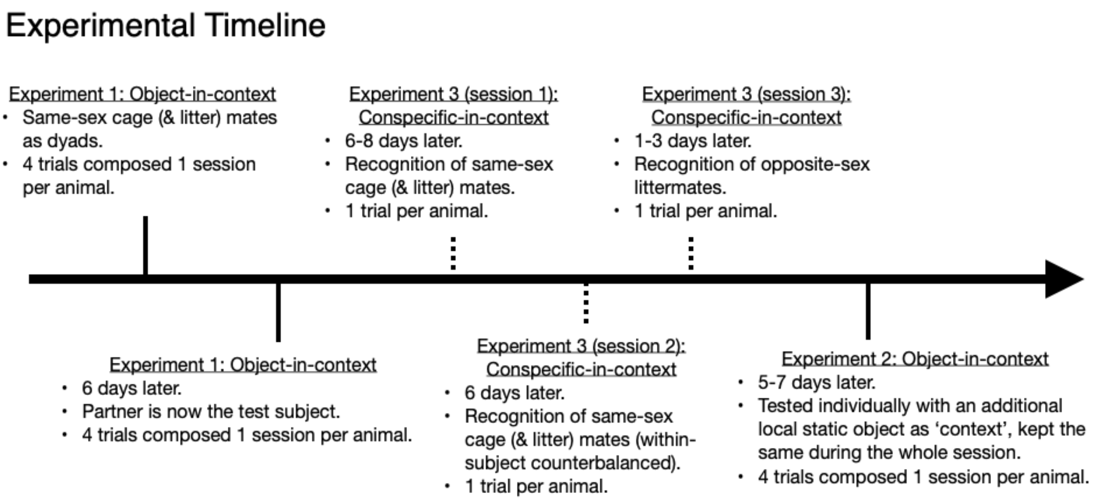


**Supplementary Figure 1.** The timeline for all the presently described experiments.

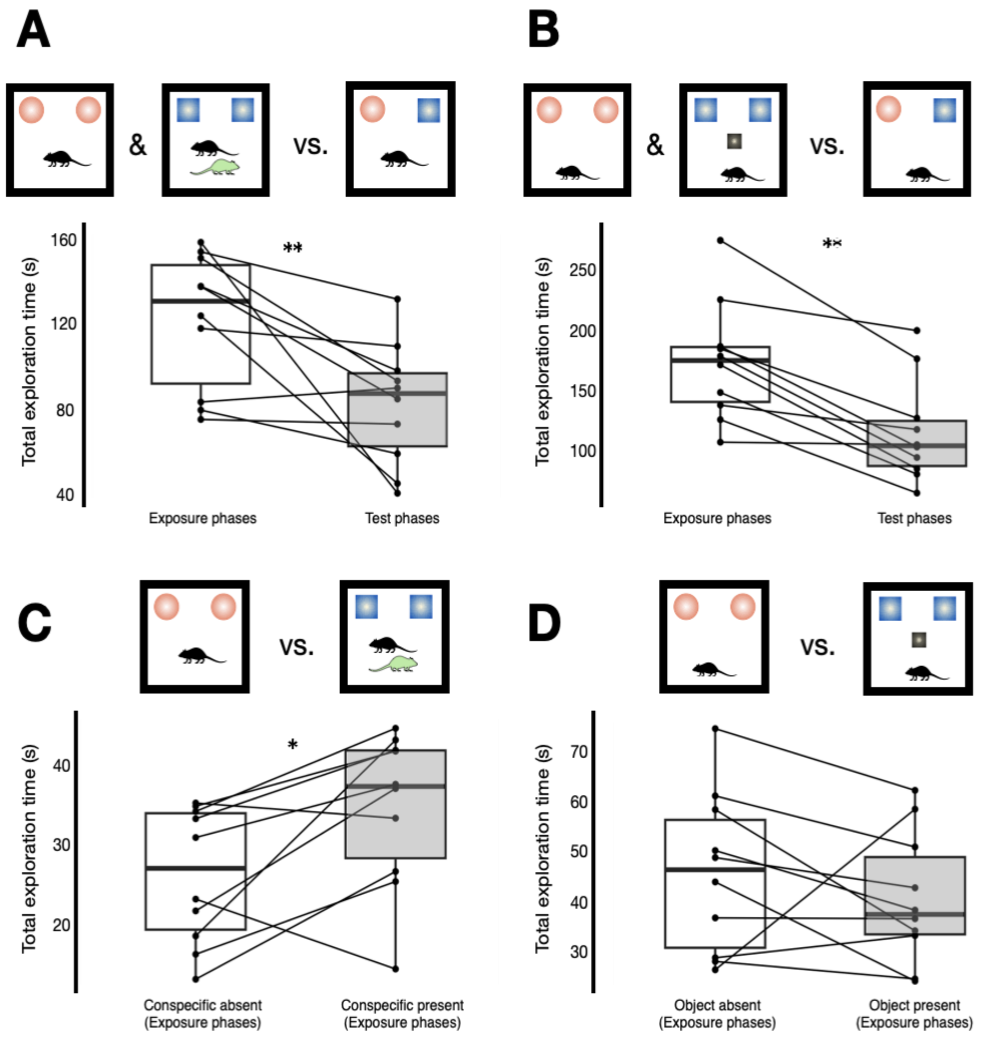


**Supplementary Figure 2.** Analyses of object exploration during the exposure phases of the object-in-context variants. (**A**) Summed total object exploration during exposure phases scaled to compare against summed total exploration in the test phases. A mixed repeated measures ANOVA yielded a significant main effect of 'context' (conspecific, experiment 1, vs. object variant, experiment 2; $F_{(1,9)} = 14.04$, $p = 0.005$, $\eta_p^2 = 0.61$). Fisher's least significant difference post-hoc (LSDph) analyses (for all following comparisons) revealed that there was significantly more exploration in Exp 2 (M = 143.71) vs. Exp 1 (M = 101.72, $p = 0.005$). There was also a significant main effect of 'phase' (exposure vs. test phase; $F_{(1,9)} = 46.70$, $p < 0.001$, $\eta_p^2 = 0.84$. Significantly more exploration in exposure phases (M = 147.17) vs. test phases (M = 98.26, $p < 0.001$). The ANOVA yielded no initial overall interaction between context and phase ($F_{(1,9)} = 1.25$, $p = 0.29$, $\eta_p^2 = 0.12$). However, post-hoc tests revealed that within context, there was significantly more exploration in the exposure phases of Exp 1 (M = 121.33) vs. the test phases (M = 82.11; $p = 0.01$, *shown in A*). (**B**) Post-hoc tests also revealed that there was significantly more exploration in the exposure phases of Exp 2 (M = 173.01) vs. the test phases (M = 114.42; $p < 0.001$). (**C**) A mixed repeated measures ANOVA was conducted for object exploration during only the exposure phases. Similarly to A, there was a significant main effect of context ($F_{(1,9)} = 12.33$, $p = 0.007$, $\eta_p^2 = 0.58$; more exploration in Exp 2, M = 43,25, vs. Exp 1, M = 30.33, $p = 0.007$). There was no main effect of 'presence' (conspecific/object presence vs. absence; $F_{(1,9)} = 0.25$, $p = 0.63$, $\eta_p^2 = 0.03$), nor 'trial-type' (test made in the 1st context vs. test made in the 2nd context trials; $F_{(1,9)} = 1.58$, $p = 0.24$, $\eta_p^2 = 0.15$). There was a significant 2-way interaction between context and presence ($F_{(1,9)} = 9.11$, $p = 0.02$, $\eta_p^2 = 0.50$). Post-hoc tests revealed that within presence, there was more exploration in Exp 2 in absence of the object (M = 45.82) vs. when mice were alone in Exp 1 (M = 26.08, $p = 0.001$). However, there was no difference between the presence exposure phases across Exp 1 and 2 (Exp 2: M = 40.96; Exp 1: M = 34.59, $p = 0.19$). *As shown in C*, within context (of Exp 1), there was significantly more exploration when there was conspecific presence (M = 34.59) vs. their absence (M = 26.08, $p = 0.017$). Finally, the ANOVA revealed no overall 3-way interaction between, context, presence and trial-type ($F_{(1,9)} = 0.11$, $p = 0.75$, $\eta_p^2 = 0.01$; within context & trial-type: test in the 1st context conspecific presence, M = 32.66 vs. alone, M = 24.64, $p = 0.09$. Test in the 2nd context conspecific presence, M = 36.52 vs. alone, M = 27.52, $p = 0.11$). (**D**) There was no difference between the exposure phases of Exp 2 (presence: M = 40.69 vs. absence: M = 45.82; $p = 0.32$). Within context and trial-type: test in the 1st context trials object acting as context present (M = 44.75), vs. absent (M = 53.53, $p = 0.43$). Test in the 2nd context trials object present (M = 36.62), vs. absent (M = 38.11, $p = 0.87$). Of note, schematics of only test in 1st context trials are shown for consistency, both trial types were considered for all the above reported analyses. *Denotes $p < 0.05$, **Denotes $p \leq 0.01$

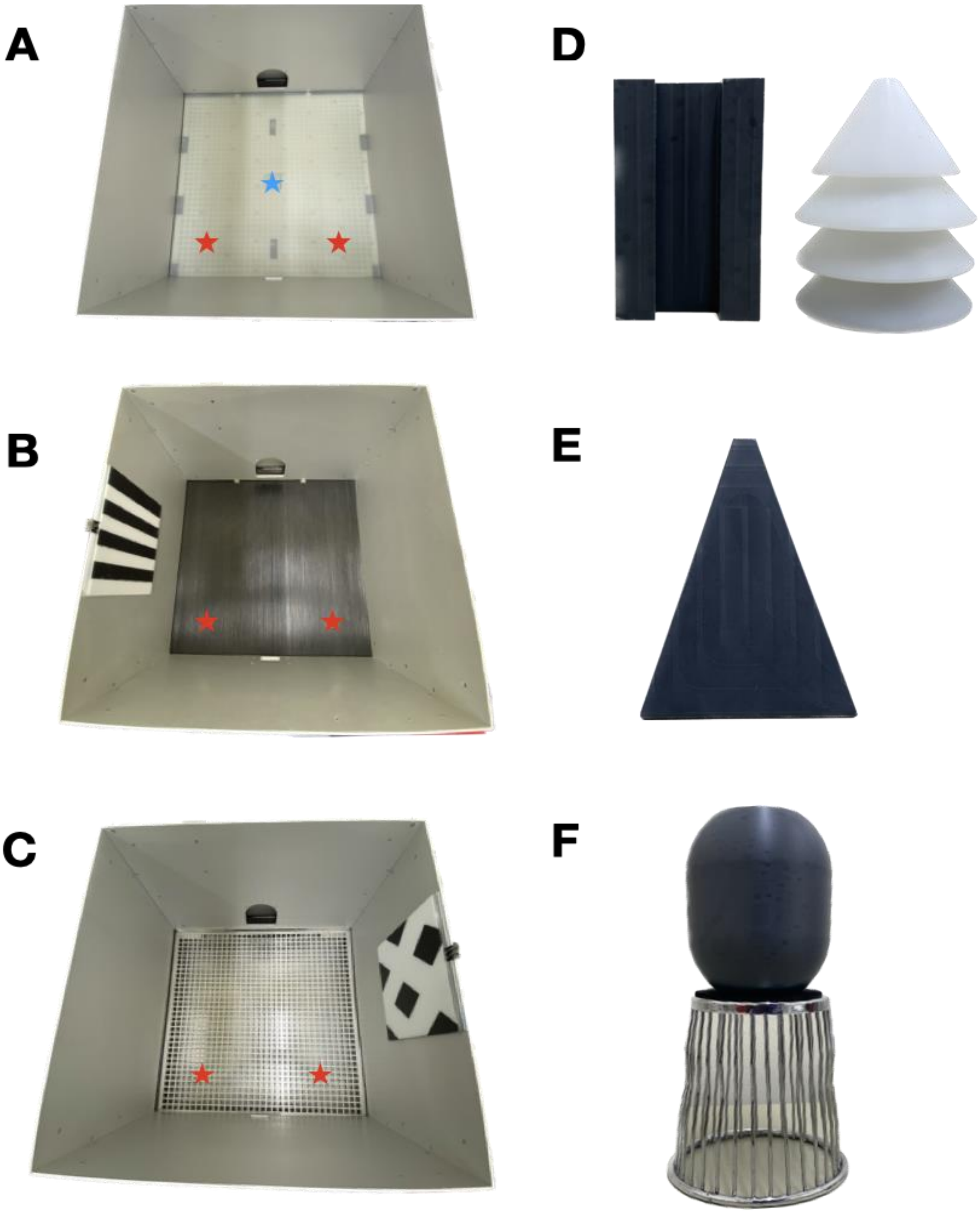


**Supplementary Figure 3.** Environment-based contexts and objects. (**A**) Context X open field, used for the object-in-context spontaneous recognition variants (experiment 1 and 2). It was comprised of no wall cues and a translucent Perspex floor with no holes. For reference, the door was considered south and the objects were placed towards the far corners north indicated via the red stars. The blue star indicates placement of the additional local object (see E) that could act as context. (**B**) Context Y open field, one of the two contexts used for the social conspecific-in-context recognition experiment 3. It was comprised of a striped, textured rubber black floor, paired with a polarised striped cue card on the east wall. Red stars indicated approximate placement of the wire cups (see F) containing conspecifics. (**C**) Context Z open field, the other context used for the conspecific-in-context social recognition experiment 3. It was comprised of steel mesh flooring paired with a polarised diamond patterned cue card on the west wall. (**D**) Example object pair used for the object-in-context experiments. Black object: 5.5 × 5.5 × 9.0cm (l × w × h). White object: 8.0cm diameter, 9.0cm height. (**E**) The additional local object acting as context, kept the same throughout the session. Position indicated via the blue star in A. It measured 5.5 × 5.5 × 7.2cm. (**F**) The chrome steel wire cup (10.2cm diameter, 10.8cm height; Model: 31570, Spectrum Diversified Designs, Inc., Ohio, U.S.A.) used to contain conspecifics, and object used to weigh it down (8.0cm diameter, 9.0cm height). See red stars in B and C for approximate placement in the environment. Of note, the lighting during experimental testing was dimmer than that depicted in A-C
